# Single-Cell and Population-Level Neuromodulation Dynamics in Dual-Electrode Intracortical Stimulation

**DOI:** 10.1101/2025.06.25.661479

**Authors:** Kayeon Kim, Victor Sonego, Beck Strohmer, Bingdong Chang, Xiyuan Liu, Alexandra Katherine Isis Yonza, Krzysztof Kucharz, Antonios Asiminas, Barbara Lind, Lechan Tao, Xiao Zhang, Shelley Fried, Anpan Han, Changsi Cai

**Author notes:** Correspondence should be addressed to: Changsi Cai, PhD., Kayeon Kim, PhD. Competing interest: No.

## Abstract

In neuroprosthetics, intracortical microstimulation (ICMS) recruits cortical networks to evoke brain responses and sensory perceptions. However, multi-electrode ICMS often generates suboptimal percepts compared to single-electrode ICMS, suggesting nonlinear neuromodulation rather than simple summation by multi-electrode ICMS. Yet, the factors and mechanisms underlying this modulation remain poorly understood. To investigate multi-electrode ICMS, we combined two-photon calcium imaging with a well-controlled dual-electrode ICMS in the mouse visual cortex to investigate how neurons integrate converging ICMS inputs at varying intensities. We found that stimulation intensity significantly shapes neuromodulation at both single-cell and population levels. Specifically, low intensities (5-7 µA) have a minimal effect on neural responses. At intermediate intensities (10-15 µA), we observed diverse, nonlinear bipolar modulation—both enhancement and attenuation—at the single-cell level. However, we achieved net enhancement at the population level. At higher intensities (15–20 µA), although the proportion of modulated neurons increased in both enhancement and attenuation directions, the net effect at the population level was neutral (zero modulation). Furthermore, neurons strongly responsive to single-electrode ICMS were more likely to be attenuated, while weaker responding cells exhibited enhanced modulation. The strongest neuromodulatory effects occur at intermediate spatial distances in between the two electrodes. Computational modeling based on spiking neural network composed of adaptive exponential integrate-and-field neurons implicated the importance of inhibitory network dynamics and network variability as key mechanisms. Our experimental data was used to train an advanced deep learning approach, which successfully predicted the neuromodulation patterns induced by dual-electrode ICMS. Our findings reveal intensity- and spatial-dependent rules of neuromodulation by ICMS, providing necessary insights to optimize multi-electrode ICMS for neuroprosthetic applications.

**Significance statement:** Understanding how cortical neurons integrate concurrent inputs from multi-electrode intracortical microstimulation (ICMS) is essential for advancing neuroprosthetic technologies. We show that dual-electrode ICMS evokes distinct, predictable neuromodulatory effects that depend on (i) stimulation intensity, (ii) a neuron’s baseline responsiveness to single electrode input, and (iii) its proximity to the electrodes. Low and intermediate intensity dual-electrode ICMS amplifies neural activity compared to single-electrode ICMS, whereas high-intensity stimulation leads to attenuation, limiting net activation.

## Introduction

Intracortical microstimulation (ICMS) significantly advances the development of brain-machine-interfaces, by providing precise, targeted stimulation within specific cortical regions. Electrical stimulation through arrays of multiple electrodes shows promise in regaining somatosensations (1–4) and for restoring visual perception through cortical and retinal implants (5, 6) in humans (7–10) and in primates (11, 12). By delivering currents through multiple electrodes, it is expected to mimic naturalistic sensation with spatiotemporal dynamics (13) or generates an integrated form of visual phosphenes for coherent shapes or letters (7, 14).

Emerging evidence suggests that distributing lower currents through multiple, closely spaced electrodes is more effective and safer than using high-intensity stimulation via fewer sites (12, 14–16). For example, a microelectrode array, such as Utah Array, a high-density microelectrode interface, can elicit cortical activity with currents as low as 4-7 µA, creating perceptually relevant stimulation patterns (11, 17). Yet, percepts induced by such multi-electrode arrays often remain rudimentary; in primates, recognition performance with multi-site ICMS is typically compromised (11, 18). This limitation suggests that cortical activation from individual electrodes may not always translate into the desired percepts when applied simultaneously, and that complex, network-level neuromodulation arising from individual neurons may shape the overall sensory outcome. These modulations may be comparable to the effect of recruiting large scale in a neuronal network, which is also known to affect the base response.

In the sensory cortex, neurons routinely integrate multiple concurrent inputs through mechanisms such as center-surround interactions in visual (19–21) and somatosensory cortices (22), lateral inhibition (23, 24), and feedback modulation from higher-order areas (25). These processes dynamically shape neuronal responses based on the spatial and temporal context of incoming stimuli, enabling flexible sensory representation and perceptual stability. As observed in natural sensory processing (19, 26, 27), spatiotemporal modulation among subpopulations of cortical neurons during artificial stimulation may affect their response selectivity and tuning beyond the direct site of stimulation. Although biomimetic stimulation protocols aim to replicate aspects of this natural integration to improve the naturalism and resolution of evoked percepts (7), how multiple artificially induced cortical inputs interact at the single-cell and network levels remains poorly understood.

While the effects of single-electrode ICMS on neuronal recruitment have been well-characterized in both excitatory and inhibitory populations (19,20), the principles governing neuronal modulation under concurrent, spatially distributed multi-electrode stimulation remains unclear. Most insights into these dynamics stem from computational models (30–32), with limited experimental evidence at cellular resolution. This raises fundamental questions which are important for brain machine interfaces (BMIs): how do neurons integrate converging inputs from multiple electrodes, and what spatial and functional features shape these interactions at both the single-cell and population levels? What are the potential underlying mechanisms?

To address these intriguing questions, we investigated single-cell and population-level neuronal responses to dual-electrode ICMS in living and transgenic mice. We combined recording excitatory neuronal calcium (Ca^2+^) responses using two-photon microscopy (TPM) with precisely controlled ICMS delivered through two closely spaced microelectrodes at varying stimulation intensities. We compared neuronal responses between single- and dual-electrode stimulation conditions and examined how neuromodulatory patterns are influenced by (i) stimulation strength, (ii) baseline responsiveness (neuronal response to single-electrode stimulation), and (iii) spatial proximity to the electrodes. Finally, computational modeling with neural network approaches was used to infer circuit-level mechanisms and assess the predictability of modulation outcomes. Our results show spatially structured, response-dependent modulation patterns during dual-electrode ICMS, offering mechanistic insights and practical strategies for optimizing multi-channel ICMS for neuroprosthetic applications.

## Results

### Distinct groups of excitatory neurons are activated by single or dual-electrode stimulation

Thy1-GCaMP6s mice expressing green fluorescent Ca^2+^ indicators in excitatory neurons were anesthetized by isoflurane and imaged by two-photon microscopy (Fig. 1A). After inserting two micro-electrodes in the exposed mouse visual cortex, we delivered either single or simultaneous electrical stimulations at varied intensities (Fig. 1B-C). The field of view (FOV) was positioned in the brain region adjacent to electrode E1 or E2 (Fig. 1D). By single-electrode E1 (sE1) stimulation, we observed spatially dispersed activation of single neurons with a higher number of activated neurons near E1 than at the distal region (Fig. 1E, sE1 stimulation). Notably, even though the second electrode (E2) was placed 558 µm from E1 in this representative mouse (Fig. 1E, sE2: single 2^nd^ electrode stimulation), it still induced robust activation of neurons within the E1 FOV (Fig. 1E, middle panel, sE2 stimulation, an example neuron in the green box).

**Figure 1.**
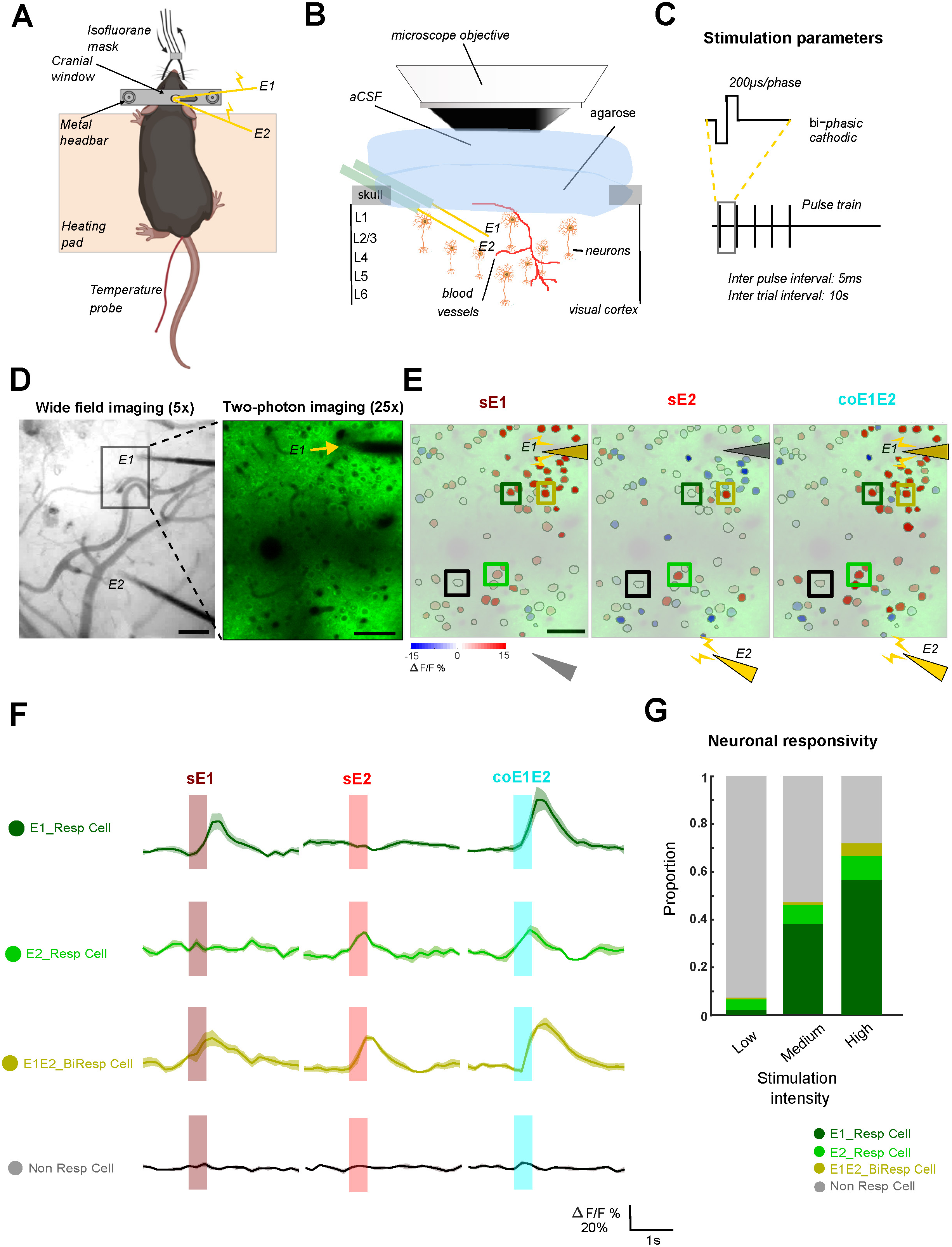
Neuronal response depends on single or dual-electrode stimulation paradigm. **A-C. Experiment scheme**. Diagrams showing electrode placement (yellow) (A), electrode placement under the microscope objective (B), and stimulation parameters (C) (see methods). **D**. **Left**: Neuronal Ca^2+^ expression under epifluorescence, scale bar, 0.1mm. **Right**: Two-photon imaging field of view (FOV), with Electrode 1 (E1) in the upper right corner, scale bar, 0.05mm. Electrode within the two-photon FOV (E1); Electrode positioned outside the FOV (E2) (see Methods). **E.** Response amplitudes mapped onto somatic ROIs across three stimulation conditions: sE1 (left): single-electrode stimulation at E1; sE2 (middle): single-electrode stimulation at E2 (outside FOV); coE1E2 (right): simultaneous (dual-electrode) stimulation at both electrodes during medium level stimulation intensity. Yellow/gray triangular arrows indicate electrode locations, with yellow as stimulation on and gray as stimulation off. Boxes show example neurons showing in (F): dark green: E1_Resp_Cell; green: E2_RespCell; Olive:E1E2_BiResp Cell; black: Non_Resp Cell. **F**. Examples of different cell response categories: E1_Resp Cell responds merely to sE1 (first row); E2_Resp Cell responds merely to sE2 (second row); E1E2_BiResp Cell responds to both sE1 and sE2 (third row). Non_Resp Cell (fourth row) shows no response to either sE1 or sE2. **G**. Proportion of responsive neurons across different stimulation intensities: Low (5-7µA, total n=486 cells, N=5 mice), Medium (10-15µA, n=557 cells, N=6 mice), and High (>15-18µA, n=507 cells, N=5 mice), pooled across all recording sites. Colors represent response categories.

Next, we asked whether each electrode activates distinct groups of neurons. The responsive neurons are categorized into four groups: ‘E1_Resp Cell’ refers to neurons that responded exclusively to sE1 (Fig. 1F, first row, dark green); ‘E2_Resp Cell’ refers to neurons that responded only to sE2 (Fig. 1F, second row, green). We also identified a subset of neurons that responded to both sE1 and sE2, referred to as ‘E1E2_BiResp Cell’ (Fig. 1F, third row, olive). Finally, ‘Non_Resp Cell’ (Non-responsive cells) did not exhibit responses to either sE1 or sE2 (Fig. 1F, bottom row, black). Although some of these ‘non-responsive cells’ exhibited increased activity during dual-electrode stimulation, we did not classify them separately from completely silent neurons due to their small population. Overall, this categorization enabled us to track how the individual neuron responses are affected by single and dual-electrode stimulation at different stimulation intensities.

We then pooled neurons across all the recording sites to quantify the number of active neurons, including both excitation and suppression from the baseline across three different stimulation intensity levels: low, medium and high (detailed definition in Methods). As expected, the proportion of activated cells increased with higher stimulation intensity (Fig. 1G). E1_Resp Cell were predominated by the sE1 electrode (Fig. 1G), as these cells were located closer to electrode E1 within the TPM FOV (Supplementary Fig. 1A) compared with E2. A substantial number of neurons in the E1 FOV also responded to sE2, and the number of sE2-activated cells increased as the stimulation intensity rose. The proportion of E1E2_BiResp cells increased from 0.61% at low intensity to 5.33% at high intensity. However, their response amplitudes to sE1 and sE2 were not significantly different (Supplementary Fig. 1B). These results suggest that sE1 and sE2 activated distinct populations of pyramidal neurons.

### Dynamic neuromodulation during dual-electrode ICMS

Next, we asked whether the responsive neurons underwent neuromodulation, i.e., enhancement or attenuation, by simultaneous dual-electrode stimulation, compared to the sum of single-electrode stimulations. Thus, we defined the neuromodulation effect as subtracting the summed responses produced by sE1 and sE2 from the response produced by coE1E2, i.e., coE1E2 – (sE1+sE2), and defined enhanced or attenuated neuromodulation as positive and negative net responses.

At the single-cell level, we observed that a substantial number of neurons were significantly modulated by coE1E2 stimulation (Fig. 2A), with some neurons showing enhanced responses (representative trace in Fig. 2B left), some exhibiting attenuated responses (Fig. 2B middle), and others showing no change (Fig. 2B right). At low and medium stimulation intensities, coE1E2 stimulation led to an overall enhancement at the population level by averaging the responses of all the detected neurons in the FOV. Interestingly, we observed net zero modulation at high stimulation intensity (Fig. 2C).

**Figure 2.**
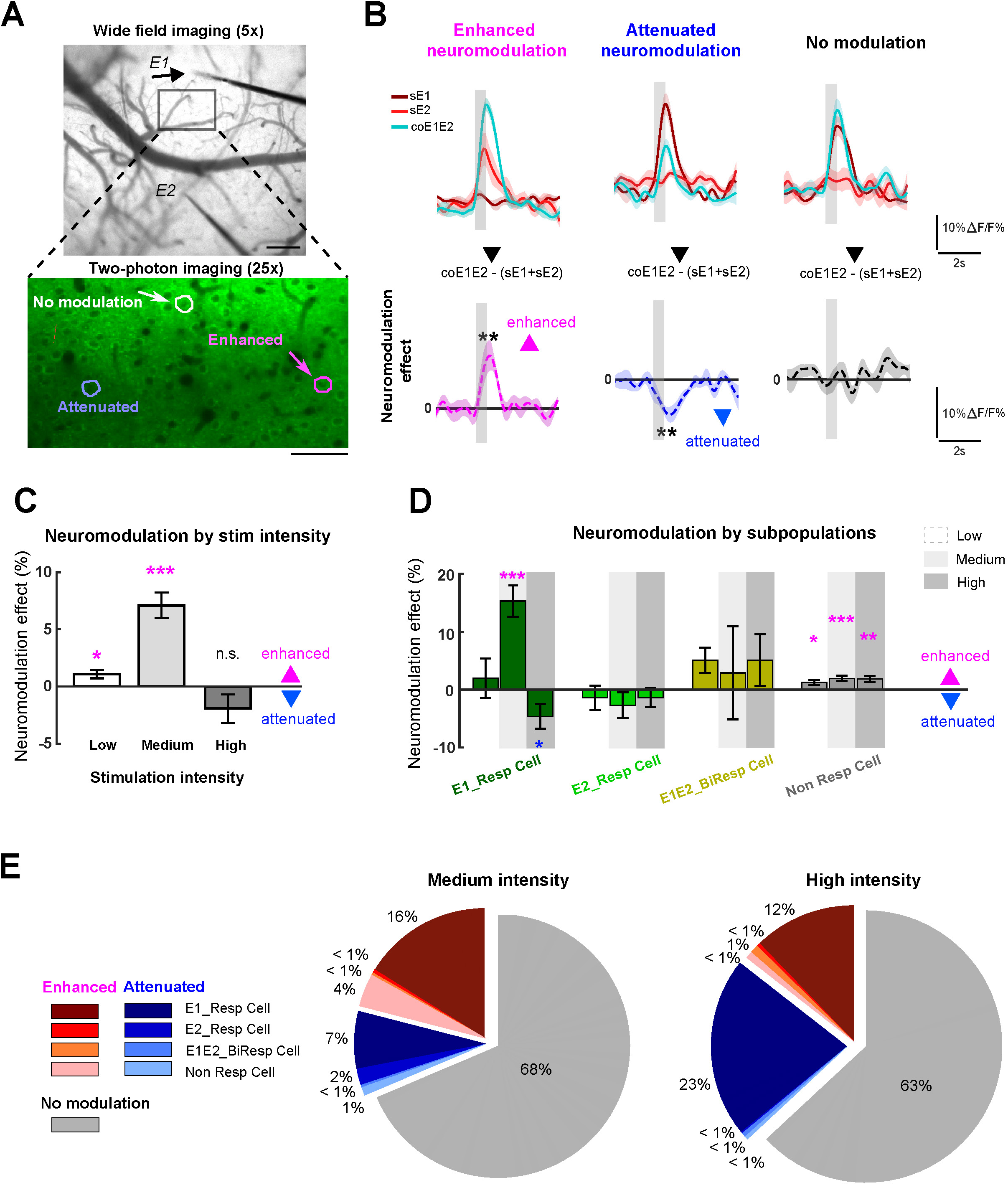
Single-cell level response modulation during coE1E2. **A**. The wide-field image (top) and TPM imaging site (bottom) are shown. Scale bar, 250 µm (wide-field), 50 µm (TPM). **B**. Example neurons with response traces averaged across 5∼10 stimulation trials (shaded area represents mean ± S.E.M.). A neuron with an enhanced effect (left) during coE1E2 compared to single-electrode stimulation (sE1 +s E2), and attenuated effect (middle), and no modulation effect (right). The locations of these cells are marked in the TPM FOV shown in A. The lower panels display the difference in response (ΔF/F %) between coE1E2 and (sE1+sE2), with significance indicated, *t*-test against 0, \*\**p*<0.001. **C-D**. **C**. Neuromodulation by population across different stimulation intensities. Mean and errorbar, ± S.E.M. **D**. Cell-type specific neuromodulation at different stimulation intensities; E1_Resp cells (dark green), E2_Resp cells (green), E1E2_BiResp cells (olive), Non_Resp cells (gray), with significance indicated, *t*-test against 0, \**p*<0.05, \*\**p*<0.005, \*\**p*<0.0001. **E**. Pie charts depicting the proportion of cells showing significant modulation (%, count) of each categorized population from all recorded neurons. Results are shown for medium (left, 557 neurons, N =6 mice) and high (right, 507 neurons, N=5 mice) stimulation intensities.

Further, we asked whether different groups of neurons (E1_Resp Cell, E2_Resp Cell, E1E2_BiResp Cell and Non_Resp Cell) demonstrated distinct neuromodulation effects by dual-electrode stimulation. At medium intensity, E1_Resp Cell showed a prominent enhancement during coE1E2 stimulation (Fig. 2D, E1_Resp Cell). However, at high intensity, the neuromodulatory effect in E1_Resp Cell dropped below zero. Since E1_Resp Cells at high intensity make up 56% of the total population (Fig. 1G), their attenuation led to an overall zero population-level modulation when neurons were pooled across all recording sites (Fig. 2C). Interestingly, across all stimulation intensities, the Non_Resp Cell group showed a significantly enhanced modulation (Fig. 2D).

To further elucidate the mechanism underlying the neutralized neuromodulatory effect at high stimulation intensity, we compared the proportions of each neuron group across the entire field of view at medium and high intensities. The proportion of neurons showing significant modulation during coE1E2 stimulation demonstrated a notable shift from enhancement to attenuation with increased stimulation intensity (Fig. 2E). Specifically, as the stimulation intensity rose, the proportion of attenuated neurons increased significantly from 10.2% at medium intensity to 24% at high intensity (χ^2^ test, *p*<0.0001). This shift was particularly pronounced in the E1_Resp Cell group. Conversely, the proportion of neurons displaying enhanced response significantly decreased from 21.3% to 14.5% (χ^2^ test, *p*<0.005). Other groups, i.e., E1E2_BiResp Cell, Non_Resp Cell, showed neuromodulation in less than 2% of population (Fig. 2E). Collectively, these results suggest higher stimulation intensity does not recruit additional responsive neurons; rather, they substantially increase the proportion of neurons attenuated during coE1E2 stimulation.

### Neuromodulation during dual-electrode ICMS depends on neuronal responses to single-electrode stimulation

We next asked what factors dominate the neuromodulatory effect of individual neurons at the single-cell level. First, we investigated whether neuronal responses to single- and dual-electrode stimulation were correlated (Fig. 3A). To examine this, we calculated the correlation coefficient between the response amplitudes to sE1 stimulation and coE1E2 stimulation for each neuron (Fig. 3B left). To assess the significance of these correlations, we compared the observed values to a surrogate distribution generated by randomly shuffling trials across the single- and dual-electrode conditions as done in our previous study (33). This analysis confirmed that the observed correlation across populations of neurons was significantly greater than chance level at the single-cell level (Fig. 3B right). At medium intensity, 32 % of neurons (68/213) showed a positive correlation between single and double-electrode responses, increasing to 41% (119/286) at high intensity. In contrast, only a small proportion showed (6/213) negative correlations (6/213 at medium; 3/286 neurons at high intensities).

**Figure 3.**
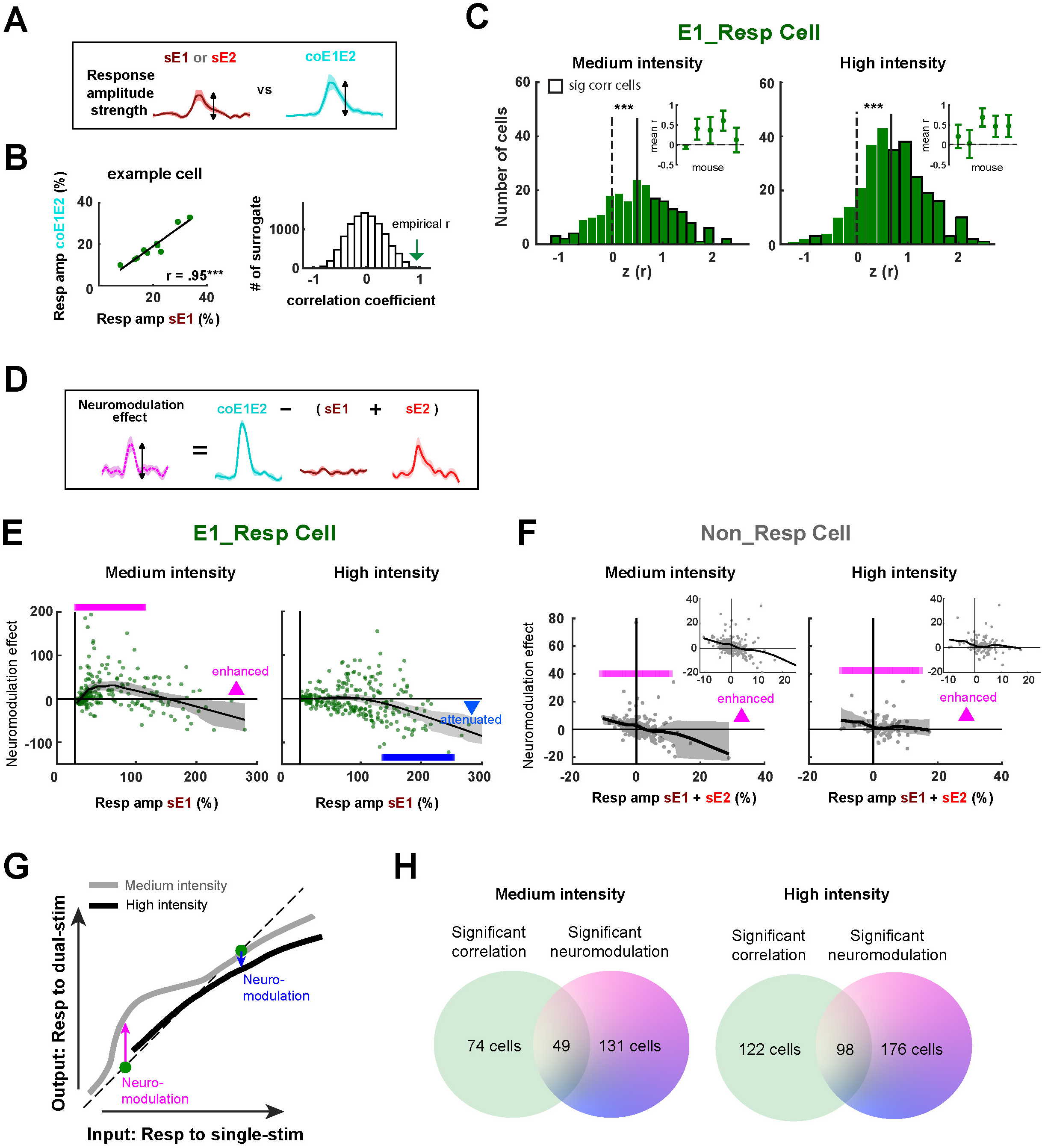
Comparison of neuronal response amplitude during single vs. dual-electrode stimulation and their relationship to neuromodulation magnitude. **(A-C).** Correlation between single- and dual-electrode responses. **A. Illustration of the quantified parameters. B.** Scatter plot of an example cell showing response amplitude (ΔF/F %) by single sE1 (x-axis) and dual-electrode coE1E2 stimulation (y-axis) (left); surrogate distribution of the correlation coefficient between single and dual-electrode response (right), with the green arrow indicating the empirical coefficient for the example cell shown on the left. **C**. Fisher-Z transformed coefficient between single and dual-electrode responses across all neurons in the E1_RespCell group. The black rectangular contour around some of the bars represents neurons with significant correlations (left: medium; right: high intensity, Monte Carlo *p*<0.05). The dashed line indicates zero, and the solid line marks the mean across all tested cells. *t*-test against 0 on Fisher-Z-transformed correlation coefficients, \*\*\**p*<0.001. Small inset indicates averaged correlation coefficient across cells in each individual mouse (N=5). (**D-F) Comparison of neuromodulation magnitude and cellular response to single-electrode stimulation. D.** Quantification of the dual-stimulation neuromodulation. **E-F.** Scatter plots showing response amplitudes during single-electrode stimulation (x-axis) and neuromodulation effects calculated as coE1E2 – (sE1 + sE2) (y-axis). Medium intensity (left panel), high intensity (right panel); **E**. E1_RespCell group; **F.** Non_Resp Cell group. Horizontal color bars indicate significance at *p*<0.05, with magenta as enhanced neuromodulation, blue as attenuated neuromodulation. The black contour denotes LOWESS fit to the data, with shaded areas indicating 95% confidence intervals by bootstrapping. Medium intensity (left panel), high intensity (right panel), *t*-test against 0, \*\*\**p*<0.001, \*\**p*<0.005. The insets are the same plots with reduced axis. **G**. An illustration of the neuromodulation pattern as a function of cell’s response to single-electrode stimulation. **H**. Pie plot illustrating the number of cells showing significant correlation, neuromodulation, and the overlap between both effects. Medium (left), high(right) intensities.

By assembling all the neurons in the E1_Resp Cell group, we found that the correlation coefficients were significantly skewed toward positive values. Similarly, positive correlations were observed in the E2_Resp Cell, Non_Resp Cell, and E1E2_BiResp Cell groups (Supplementary Fig. 2).

We next investigated whether the neuromodulatory effect (Fig. 3D) at the single-cell level is associated with the neuronal base responsiveness, defined as the neuronal responses to single-electrode stimulation. Our analysis revealed a distinct pattern: at medium intensity, coE1E2 stimulation significantly enhanced neuromodulation in neurons with the base responsiveness below 110% (ΔF/F) (Fig. 3E, left, *p*<0.05). Conversely, neurons demonstrating stronger base responsiveness (≥110% ΔF/F) showed no significant neuromodulation under these conditions. When assessing neuromodulation at high intensity by sE1 stimulation, the neurons with high base responsiveness (≥110% ΔF/F) exhibited attenuated modulation (Fig. 3E, right, E1_RespCell, *p*< 0.05). This effect could result from detector or image processing saturation at high ΔF/F values, but this explanation is unlikely to fully account for the findings because we observed both increases and decreases in activity across different neurons within the same field of view (Fig. 2A, B), indicating that our imaging maintained sufficient dynamic range to capture bidirectional modulation.. Neurons categorized as weak and moderate responders (base responsiveness < 110% ΔF/F) did not show significant modulation. Furthermore, E2_Resp Cell group did not show a clear relationship with base responsivity (Supplementary Fig.3), likely due to the limited size of this neuronal group. However, the Non_Resp Cell group demonstrated a pronounced association, with all these neurons showing enhanced modulation at both medium and high intensities (Fig. 3F).

Collectively, these results indicate that neuromodulation induced by coE1E2 stimulation is intricately associated with both a neuron’s base responsiveness and the intensity of electrode stimulation. At medium stimulation intensities, dual-electrode stimulation enhances modulation primarily in neurons with lower to moderate base responses, whereas at high stimulation intensities, modulation becomes attenuated in neurons with strong base responses. Figure 3G summarizes these observations, illustrating how neuromodulatory patterns vary as a function of each neuron’s baseline response level, thereby highlighting the nonlinear input-dependent nature of neuromodulation under this context.

To further explore the relationship between baseline responsiveness and modulation, we examined the proportion of significantly modulated cells whose responses correlated with those to single-electrode stimulation. Interestingly, the majority of dual electrode modulated neurons (73% at medium intensity and 64% at high intensity) did not depend on their single-electrode responses (Figure 3H). This further indicates that neuromodulatory effect cannot be explained by correlation between single and dual-electrode stimulations, which we discuss in detail later.

### Neuromodulation peaks between the two stimulating electrodes, and diverges with increasing stimulation intensity

So far, our results underscore that neuromodulation induced by dual-electrode stimulation depends significantly on the neuronal base responsiveness and stimulus intensity. This raises the subsequent question of the spatial localization of neurons displaying enhanced or attenuated modulation: Is there a distinct spatial distribution or an optimal region exhibiting pronounced neuromodulatory effects? Given the wide variation in neuronal response amplitudes during single-electrode stimulation, physical factors, particularly the distance between the neuronal soma and stimulating electrode, are likely influential. This is consistent with the established understanding that the strength of electric fields decreases with increasing distance, affecting neuronal responsiveness (34). Therefore, we hypothesize that this soma-to-electrode distance critically modulates the neuromodulation magnitude, determining enhancement or attenuation effects (Fig. 4A, E).

**Figure 4.**
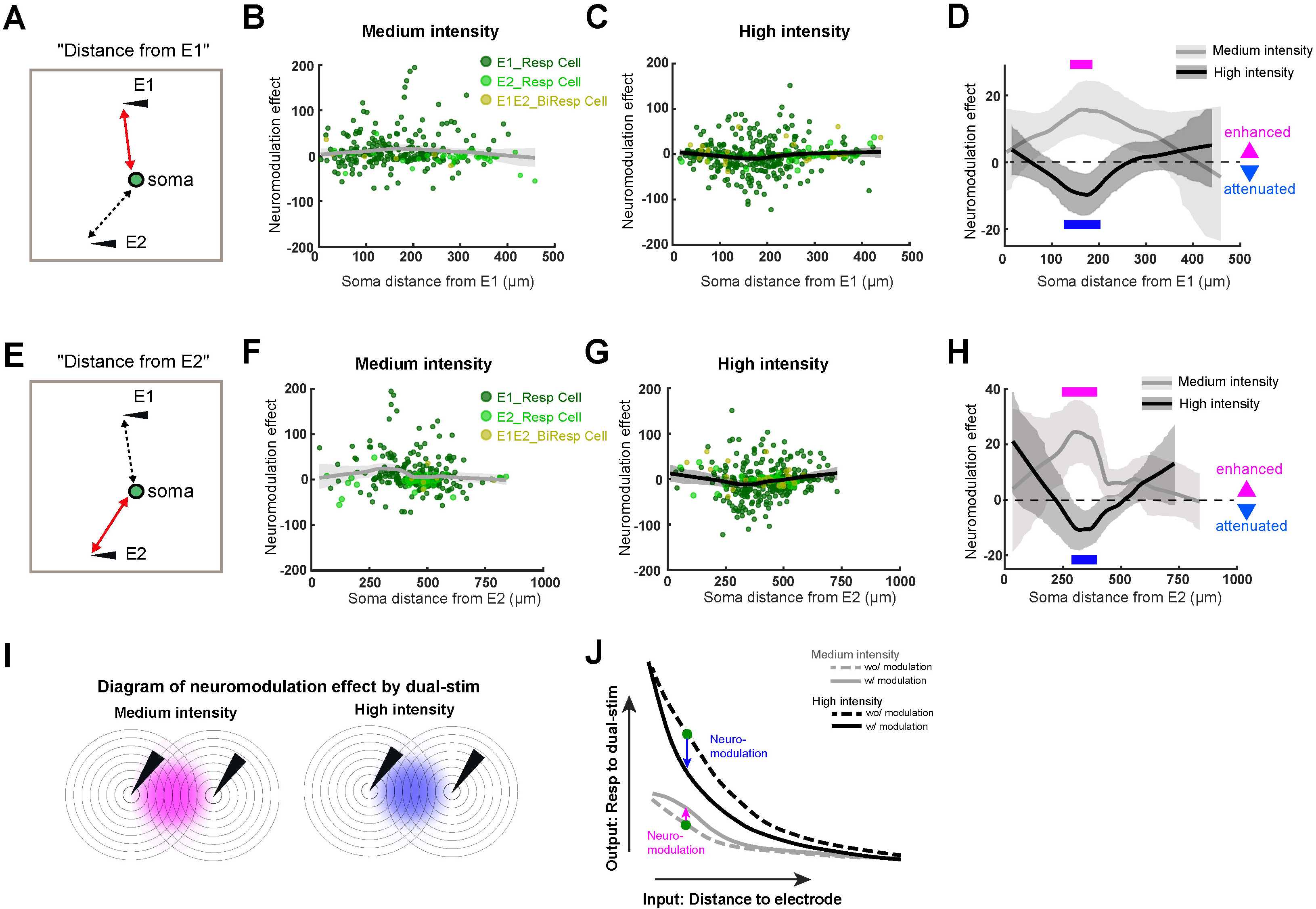
Diverging soma-electrode-distance-associated modulation effects by different stimulation intensities. **A.** Distance metric between the soma and E1 location. **B-D**. Modulation effect (ΔF/F %) as a function of soma-to-electrodes distance (µm) at medium intensity (B) and high intensity (C). Colored dots represent individual cells from E1_RespCell (dark green), E2_RespCell (green), and E1E2_BiRespCell (olive) groups. **D**. Fittings from B and C overlaid for medium (gray) and high (black) intensities. Horizontal lines indicate significance based on Monte-Carlo *p*<0.001, FDR corrected (see Methods). **E-H**. Same convention as A-D, but showing soma distance relative to the E2 location (E). **I, J**. Diagram of the neuromodulation effect by dual-electrode stimulation at medium (I, left) and high (I, right) intensities. Illustration of the response to dual-electrode stimulation as a function of the distance between the cell and the stimulating electrode (J).

Using a LOWESS fitting curve (35), we found neuromodulatory enhancement at 156∼178 µm from E1 at medium intensity at the population level (Fig. 4B, D). At high intensity, neuromodulatory attenuation was observed at 126-204 µm from E1 (Fig. 4C-D). These results indicate a marked location divergence, where modulation effects occur at approximately the same distance from the two electrodes, but with different outcomes depending on the stimulation intensity (Fig. 4D, Monte-Carlo *p* <0.05, FDR corrected). A similar divergence pattern was observed when somata locations were aligned based on their distance from E2 (Fig. 4E-H), though with a slight positional shift of the modulation peaks. This spatial dependency is further illustrated in Fig. 4I-J, which summarizes the strong relationship between neuromodulation effects and the distance between the soma and the stimulating electrode. Notably, this diverging pattern between different stimulation intensities was absent in Non_Resp cells (Supplementary Fig. 4A-B).

### The modulation effect is attributed to inhibitory and noise inputs during dual-electrode stimulation

It is well characterized that single-electrode ICMS recruits both excitatory and inhibitory neurons, with increasing stimulation intensity particularly stimulating inhibitory neurons that are located further from the stimulating electrode (29). Although we only examined the responses of excitatory pyramidal neurons, our results showing dominant attenuation, especially between the electrodes during high intensity dual-electrode ICMS, indicate the potential recruitment of inhibitory neurons.

To gain deeper insight into the possible circuit mechanisms underlying modulation effects in a dual-electrode stimulation setup, we developed a computational model using a spiking neural network composed of adaptive exponential integrate-and-field neurons (Eq 1,2,(36)). The network architecture consisted of excitatory (E) and inhibitory (I) populations, following the connectivity structure outlined by O’Rawe et al. (37). Stimulation was delivered via two electrodes (Fig. 5A), with randomized soma locations (Fig. 5B), as described in the Methods section. Figure 5C (upper) visualizes the comparison of the location versus the effect of stimulation on each neuron in a sample trial. The plot illustrates that the general trend of enhancement and attenuation effects tend to be stronger near the midpoint. The accompanying model include both control conditions and various experiment perturbations.

**Figure 5.**
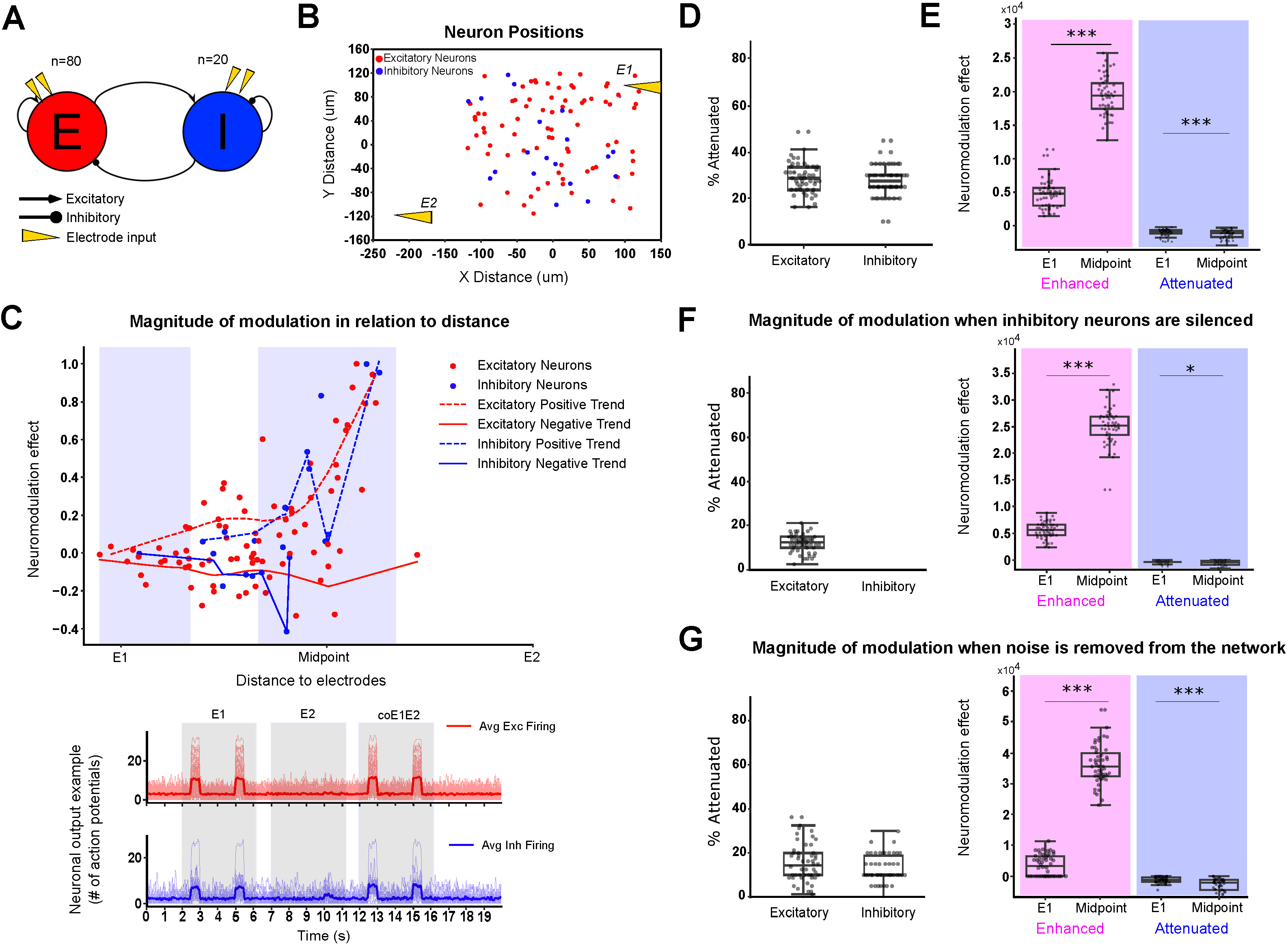
Computational modeling. **A-B.** Visualization of the computational model representing visual cortex. **A.** Block diagram showing the connectivity between populations, excitatory (E) neurons excite the inhibitory (I) neurons while the inhibitory neurons inhibit the excitatory neurons. Each population is also self-connected. The dual-electrode stimulation is represented with the orange triangles. The populations are stimulated by the same electrodes. **B**. Sample scatter plot representing the neurons and electrodes in space, point (0,0) represents the center of the neuronal population. **C**. (Upper): Sample modulation from a single test showing the amount of enhancement or attenuation of a neuron in relation to location, the magnitude of modulation (=modulation effect) is determined by comparing area under the curve (AUC) during current pulses. Blue shaded areas show which neurons are considered “close” to E1 or the midpoint; (Lower): Sample output of individual neurons in a single trial. Neuron firing is represented by binning action potentials using a 5ms time window. The dark trace is the average of all neurons of that type (red=excitatory, blue= inhibitory). The electrodes provide current at 2 second intervals, for a period of 500ms, meaning a complete on/off cycle is 2500ms. The first two current pulses are from E1, the next two from E2, and the final two are from both E1 and E2. **D-E**. Control results showing modulation of neural activity by population (n=50). **D**. Variance plot showing the percentage of neurons attenuating under dual electrode stimulation. **E**. Variance plot showing the amount of enhancement and attenuation of excitatory neurons located closer to E1 versus close to the midpoint. **F**. Same convention as D and E, but without inhibitory input. **G**. Same convention as D and E but without noise input. Wilcoxon Signed-Rank test, \*\*\**p*<0.001, \*\**p*<0.01, \**p*<0.05.

First, our connectivity model shows a mean percentage of attenuated excitatory neurons of 28.8±6.43% and 27.8±6.18 % (Fig. 5D) of inhibitory neurons. The proportion of attenuated neurons falls close to the range observed at medium to high stimulation intensities in our empirical data (Fig. 2E). To quantify modulation, we analyzed neuronal output traces (Fig. 5C, lower) and calculated response enhancement and attenuation across dual-pulse (coE1E2) and single pulse stimulation trials (sE1+sE2) by comparing the area under the curve when electrode stimulation was active.

Notably, neurons positioned closer to the midpoint between the electrodes experienced stronger enhancement and attenuation effects (Fig. 5E) (Enhanced E1: 4753.66±2039.83, Attenuated E1: −938.63±464.95; Enhanced midpoint: 19426.71±2039.83, Attenuated midpoint: −1245.71±682.64), demonstrating a spatially dependent modulation pattern (Fig. 5E, upper) that closely mirrored our experimental findings (Fig. 4). Our model, which incorporates distance-dependent connectivity (Supplementary Fig. 5), stochastic noise, and spatially randomized neuron locations, successfully replicates key neuronal response trends observed in our empirical data. Namely, an increase in the modulation of neurons close to the midpoint between electrodes and a stronger response enhancement as compared to attenuation.

To further dissect the role of inhibition in neuronal modulation during dual-electrode stimulation, we examined two experimental scenarios: (1) Does silencing inhibitory neurons amplify modulation effects? (2) Given that noise is associated with both inhibitory influence (38) and information processing (39), how does removing noise input affect modulation?

As expected, silencing inhibitory neurons – by eliminating their connectivity both from the electrodes and within the network - reduced the percentage of attenuating excitatory neurons to 12.05±4.04% and eliminated attenuation in inhibitory neurons since these cells were silenced (Fig. 5F, left). This also led to a substantial increase in excitatory neuron enhancement near the stimulation midpoint (Fig. 5F, right, 25075.04±3397.05), significantly higher than control (Fig. 5E, 19426.71±2039.83, *p*=1.78×10^-15^).

Similarly, removing noise from the network also reduced the percentage of attenuating excitatory neurons to 15.28±8.38% and decreased attenuating inhibitory neurons to 12.30±6.87% (Fig. 5G, left). Notably, the enhancement of excitatory neurons near the midpoint increased even further under no-noise condition (Fig. 5G, right, 36055.54±6161.23), which was significantly higher than both control (Fig. 5E, *p*=1.78×10^-15^) and the no-inhibition condition (Fig. 5F, *p*=5.86×10^-14^). In contrast, enhancement at E1 was lower (: 3493.15±3401.46) than in the no-inhibition condition, while attenuation at both E1 (− 1196.78±940.76) and the midpoint (−3218.20±2962.42) remained present.

Collectively, our computational modeling indicates that the pronounced neuromodulation effect at the middle point of two electrodes is possibly attributed to the recruitment of inhibitory interneurons.

### Deep learning model effectively predicts neuronal responses during dual-electrode stimulation

With the expanding role of deep learning, particularly of neural networks, in neuroprosthetic technologies and brain-computer interfaces (BCIs) (40, 41) we next asked whether a deep learning model could successfully learn the spatial and stimulation-related features of the neuronal Ca^2+^ responses, and further could reliably predict neuromodulation during dual-electrode stimulation. For this purpose, we employed a Multi-Layer Perceptron (MLP) Regressor – a feedforward artificial neural network designed to capture complex, nonlinear relationships between input features and continuous outputs (42, 43), such as the average response of individual neurons.

The model was trained using empirical neural data collected across all stimulation conditions (Fig. 6A, B), with input features including geometry, electric field and stimulation intensities (Fig. 6C, Table 2). The network architecture comprised three hidden layers with 100 units each (Fig. 6E, details in the Methods). The training loss curve demonstrated reliable convergence with a low final loss (Supplementary Fig. 6A), while the learning curve confirmed strong generalization with minimal overfitting (Supplementary Fig. 6B).

**Figure 6.**
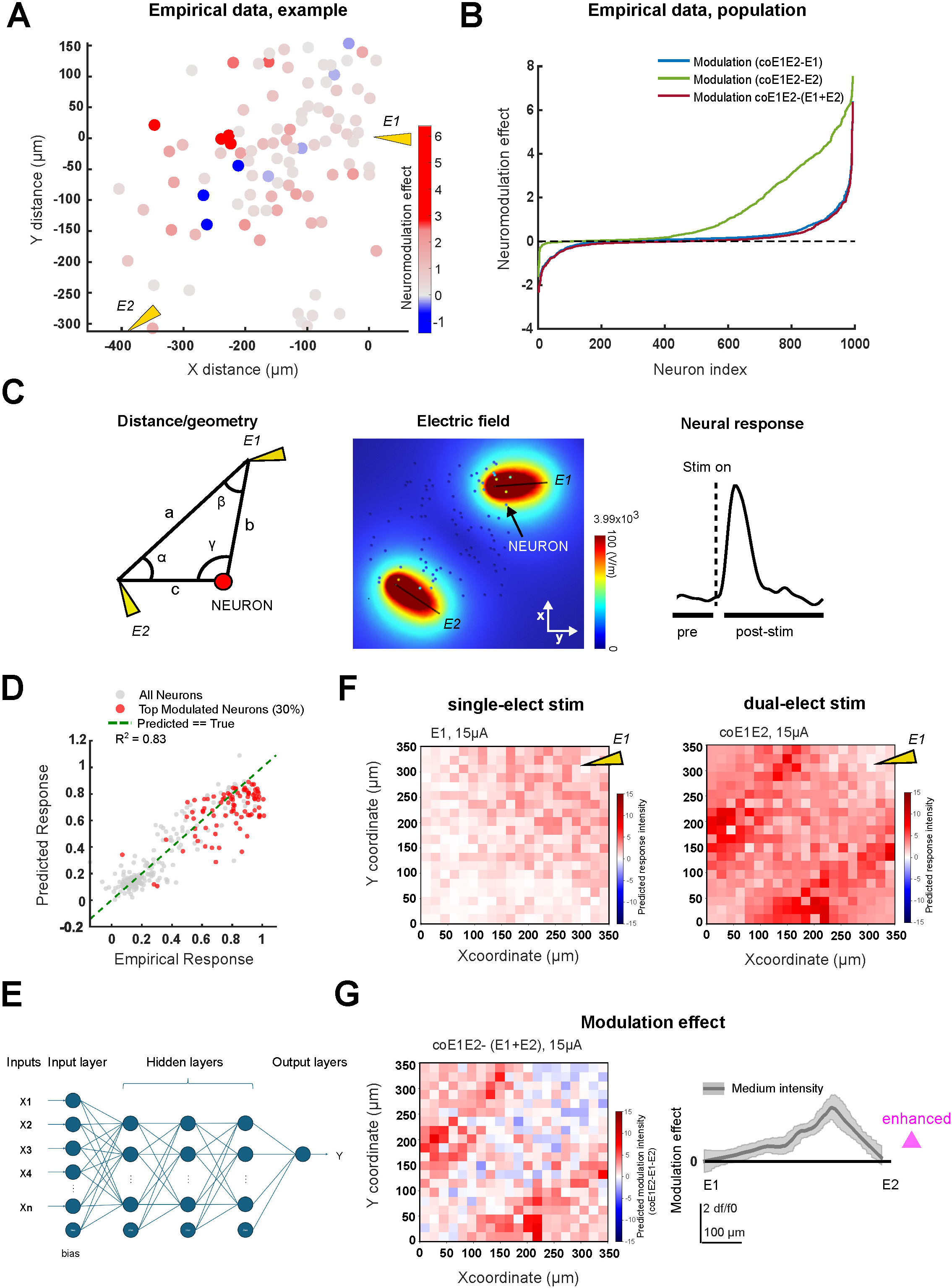
Neural network modeling. **A.** Neuromodulation plot showing the distribution of neuromodulation in space in a representative case, defined as coE1E2 – (sE1+sE2). **B.** Sorted neuronal response modulation (N = 994 neurons), difference between coE1E2 from sE1 (blue), sE2 (green), and sE1+sE2 (red). **C.** Input features and target variable: electrode and soma distance/angles (left), electric field modeling derived by COMSOL (middle), and pre-vs. post-stimulation activity (right). **D.** Scatterplot of predicted vs. true neuronal responses, with red dots marking highly modulated neurons. **E.** MLP regressor with three hidden layers (100 neurons each), ReLU activation, bias term, and scaled input features. Optimized with Adam, trained for up to 1000 iterations, α = 0.0001. **F**. Spatial map of simulated response intensity during E1 stimulation alone (left) and coE1E2 stimulation (right). Each grid cell represents the location of a single neuron. The yellow arrow indicates the position of the E1 electrode. **G**. Same convention as F, but showing the modulation effect of the predicted responses, calculated as ‘coE1E2-(sE1+sE2)’ for each neuron showing as a spatial map (left). The modulation effect was averaged along the soma’s distance from the E1 electrode location and LOWESS curve was fitted to the data, with the 95% confidence interval shaded in gray (right).

**Table. 1.**
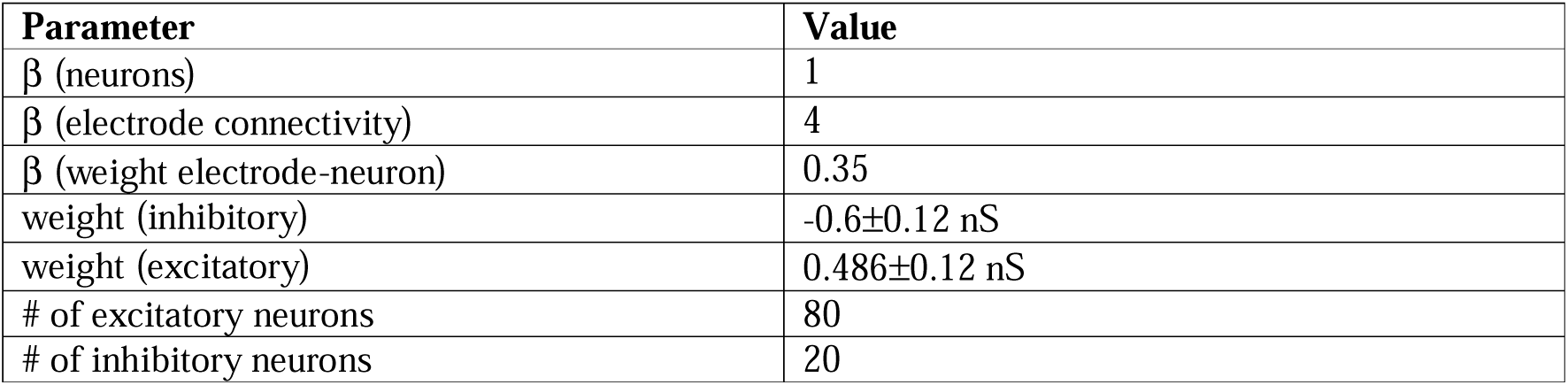
Network parameters for computational modeling.

| Parameter | Value |
| --- | --- |
| $\beta$ (neurons) | 1 |
| $\beta$ (electrode connectivity) | 4 |
| $\beta$ (weight electrode-neuron) | 0.35 |
| weight (inhibitory) | -0.6±0.12 nS |
| weight (excitatory) | 0.486±0.12 nS |
| # of excitatory neurons | 80 |
| # of inhibitory neurons | 20 |

**Table 2.** MLP Regressor input features.

| Features | Description | Unit |
| --- | --- | --- |
| Spatial | Distance, angle from neuron to E1 and E2 (5 features) | $\mu\text{m}$, $^\circ$ |
| Electrode geometry | Distance between electrodes | $\mu\text{m}$ |
| Stimulation parameters | Current intensity from E1 and E2 (2 features) | $\mu\text{A}$ |
| Pre-stimulation activity | Range of $\Delta F/F_0$ activity before first stimulation start | $\Delta F/F_0$-normalized average pixel intensity |
| Electric field | Norm of field magnitude at soma (COMSOL-simulated) | V/m |
| Interaction terms | 7 additional features computed from pairwise interactions | Various |

As a result, our MLP neural network model achieved high predictive performance, with a maximum R^2^ of 0.83 and a mean squared error (MSE) as low as 0.02 on predicted responses compared with empirical response (Fig. 6D). In Figure 6D, the red dots indicate the top 30% most strongly modulated neurons, indicating the model’s capability to predict the response of neurons that are highly modulated during dual-electrode stimulation. Next, we developed a grid of 20×20 phantom neurons spaced 17.5µm apart, for a total field of view of 350µm^2^. Our deep learning model was used to predict the responses of the simulated neurons to single-(Fig. 6F, left) and dual-electrode stimulations (Fig. 6F, right). Notably, a similar pattern of enhanced neuromodulation in the middle point between electrode E1 and E2 was well recapitulated by the model (Fig. 6G). The spatial distribution of modulation effects (Fig. 6G, left) showed a dispersed pattern between the two electrodes, with the strongest modulation occurring in the region between them (Fig. 6G, right).

These findings support the idea that dual-electrode neuromodulation produces a structured, learnable pattern of single-cell modulation that can be reliably predicted from spatial and stimulation features alone, and pioneer in using a deep learning model to study the neuromodulation effect by multi-electrode ICMS.

## Discussion

This study addresses fundamental questions on how cortical neurons integrate converging inputs from dual electrical stimulations sites, and what spatial and functional properties influence these interactions at both single-cell and network level. By combining TPM with concurrent dual-electrode ICMS, we found that distinct neuronal populations were recruited by individual electrodes, with limited overlap in those activated by the second electrode. Within these populations, up to 40% of these neurons exhibited neuromodulation when inputs converged: at low and medium intensities, population activity was significantly enhanced, while high intensity produced zero net modulation due to increased attenuation of single-cell level responses. Importantly, neurons that responded weakly to single-electrode stimulation were preferentially enhanced by dual-electrode ICMS, whereas strong responders were more likely to be attenuated. The strongest modulation effect was present in a distance-specific manner, diverging with stimulation intensity. Our computational simulations support that inhibitory network influence the modulatory effects, and a deep learning-based predictive model accurately captured the spatial and response-dependent modulation patterns induced by dual-electrode ICMS. Together, these results demonstrate that converging electrical inputs engage cortical networks through both spatially and response-dependent mechanisms, giving rise to structured, predictable patterns at the level of individual neurons.

### Synergistic recruitment and response amplification with dual-electrode ICMS

In our experiments using Thy1-GCaMP6s mice under anesthesia, increasing single-electrode current activated progressively larger fractions of excitatory neurons. This is consistent with prior reports in anesthetized mice (28, 44–46), awake primates (47), and both excitatory and inhibitory neurons in awake mice (29). Importantly, dual-electrode ICMS not only preserved this activation pattern but, in many cases, amplified single-cell and population-level responses. Specifically, dual-electrode ICMS enhanced population-level neuronal responses from the group of neurons that were not activated by single-electrode stimulation, likely due to overlapping activation fields between the two electrodes (48). These overlapping fields can be extrapolated to the other ICMS applications, because our two electrodes were located at a comparable distance to the other commercially available multi-electrode arrays for clinical and pre-clinical studies (11, 49, 50), and our stimulation intensities were also in a comparable safety range (16, 51). Particularly, this modulation effect was most pronounced in neurons located around these overlapping areas, where medium stimulation currents induced enhancement and high currents led to attenuation. These results provide strong cellular-level evidence identifying an optimal spatial zone for driving neuromodulatory effects during dual-or multi-site ICMS (52–54). Together, our data demonstrates that dual-electrode ICMS can generate synergistic interactions, enhancing cortical responses at the population level. This supports the efficacy of distributed, multi-electrode stimulation strategies for enhancing cortical activation (2, 14, 18, 55).

### Non-random, response-dependent neuromodulation

Individual neurons appear to be tuned to their dominant input (sE1, based on the FOV), which establishes their baseline responsive state. When this dominant input is mild, dual-electrode ICMS enhanced the responses of weak or moderate responders by integrating secondary input (E2) at medium intensity. In contrast, when the dominant input was strong, additional neurons with higher response amplitudes were recruited, and the secondary input tended to attenuate responses. These patterns suggest that dual-electrode neuromodulation may engage a partially structured, functionally organized, but nonlinear integration of converging inputs at the single-cell level (Fig. 3G).

Importantly, these neuromodulatory effects during dual-electrode ICMS were not solely determined by a neuron’s spatial proximity to the stimulating electrodes. A neuron’s response to single-electrode stimulation strongly predicted how it would be modulated under dual-electrode ICMS. This response-dependent modulation pattern points to a non-random, structured integration of inputs within cortical networks. Although the precise cellular and network mechanisms remain to be identified, possible contributors include differences in intrinsic excitability (56), synaptic integration properties (35), and local networks connectivity (57–62). Moreover, while the specific intrinsic and network variables underlying these differences could not be directly determined in this study, the relationship between single-electrode responses and dual-electrode ICMS modulation proved highly predictable.

Our machine learning model, trained on stimulation parameters and geometric features reflecting the location of neurons relative to electrodes, successfully predicted modulation outcomes, including neurons exhibiting the strongest effects. Notably, this was achieved even though the variability in empirical dual-electrode responses could not be fully explained by single-electrode response variability (Fig. 3C), or the magnitude of response modulation alone (Fig. 3E-F). This suggests that additional, complex relationships between the stimulation parameters contribute to neuromodulatory outcomes.

### Implications for neuroprosthesis and perceptual restoration

These findings carry important implications for the design of neuroprostheses. Rather than relying on currents delivered through a single electrode to surpass perceptual thresholds, distributing moderate currents across multiple electrodes may more effectively recruit cortical circuits while reducing suppression and minimizing off-target effects (63). This strategy could perhaps enhance the selective engagement of functional subnetworks – such as orientation columns in the visual cortex – with potential applications ranging from basic animal research (64) to sensory restoration in non-human primates (11), and human clinical settings (63, 65, 66). Prior psychophysical studies have similarly shown that multi-electrode, lower-current strategies improve perceptual resolution and detection thresholds compared to conventional single-electrode stimulation methods (15).

However, our results also reveal important limitations to this approach. Highly modulated regions (Fig. 4J), including areas exhibiting suppressive effects that extend over broader cortical distances (>300µm, roughly the spacing between E1 and E2), were frequently observed among neurons strongly responsive to either electrode during dual-electrode ICMS at higher intensities. Such widespread suppression could paradoxically reduce or negate perceptual benefits, especially when attempting to evoke fine, focal percepts like isolated phosphenes in visual prostheses (11). These findings suggest the dual nature of neuromodulation – capable of both enhancing and suppressing neural activity depending on stimulation intensity, electrode configuration, and baseline neuronal responsiveness. Therefore, careful optimization of stimulation parameters, including electrode placement, current amplitude, and targeting of responsive neural populations, will be essential to maximize the efficacy of multi-electrode or furthermore, multi-site stimulation strategies in both experimental and clinical neuroprosthetic applications.

## Acknowledgements and funding sources

This paper was supported by the Lundbeck Foundation (#R402-2022-1530, #R345-2020-1440, #R436-2023-1125, #R392-2018-2266 and #R345-2020-1440), the Independent Research Fund Denmark (#1133-00016B and #1030-00374A), the Novo Nordisk Foundation (#0064289, #0092323, #117272 and #14CC0001), Læge Sofus Carl Emil Friis og Hustru Olga Doris Friis’ Legat, Dagmar Marshalls Fond, Familien Hede Nielsens Fond and Hørslev fonden, and Oda og Hans Svenningsens Fond. An animal image was created using Biorender.

## Data sharing plans

Data and codes for figure plotting, network, and machine learning will be publicly accessible.

## Author contributions

**Conceptualization**: C.C, S.F, K.Kim; **Methodology**: C.C, K.Kim, V.S, B.S, B.C, X.L; **Investigation**: K.Kim, V.S, B.S, A.Y; **Data Analysis**: K.Kim, V.S, B.S,B.C; **Visualization**: K.K, C.C, B.S, K.Kim; **Resources**: C.C, A.A, A.H, X.Z, L.T; **Funding acquisition**: C.C, A.H; **Writing-original draft**: K.Kim, C.C, V.S,B.S; **Writing – review & edit**: all authors.

## Materials and Methods

### Animals

We used Thy1-GCaMP6s (67) mice over 8 weeks old (N=7, including both males and females) with a weight range of 20∼35g. The study followed the guidelines of Directive 2010/63/EU of the European Parliament and Council on the care and use of animals for research. All procedures received approval from the Danish National Committee on Health Research and in accordance with the European Council’s Convention for the Protection of Vertebrate Animals used for experimental and other scientific purposes, adhering to established animal research protocols.

### Experimental procedures

#### Animal preparation

During the acute experiment, 4% isoflurane was used to induce anesthesia, and 0.9-1.5% was used to maintain it throughout the experiment. After isoflurane induction, lidocaine (10mg/kg) was administered for local anesthesia before making the surgical incision. Following the craniotomy, the dura was removed. After inserting two electrodes, agarose was applied to stabilize both the cortical surface and the electrodes. Body temperature was maintained at 37 °C throughout the procedure using a heating pad.

#### Wide-field and two-photon imaging

For wide-field imaging, we used Olympus U-HGLGPS light illumination system and a UPlanSApo 4x objective with an 0.16 numerical aperture. Each pixel corresponded to 1.81µm, and the field of view measured 1440 x 1920 pixels. The vasculature captured from wide-field imaging was used as a reference point for precisely locating the electrodes during two-photon imaging and analysis. We used a two-photon microscope (FluoView FVMPE-RS, Olympus) equipped with a femtosecond laser (Mai-Tai DeepSee), using 25L×L1.05 NA water-immersion objective, as well as GaAsP detectors (68). The excitation wavelength was 920nm, and image acquisition frequencies ranged from 1.22∼2.44 Hz.

#### Stimulator and experimental protocol

We used ISO-flex (A.M.P.I.) connected to Platinum-Iridium micro-electrodes (8-11kΩ, tip diameter, 2-3µm; PI2PT30.01. A3; Microprobes for Life Sciences). The reference electrode was positioned under the neck skin of the mouse, with separate connections for each of the two electrodes. Each electrode was secured to a micromanipulator arm separately (Fig. 1A), and inserted at an angle of 15-20° for two photon imaging. The insertion was 200µm from the cortical surface, which served as the designated stimulation depth.

Stimulation was applied using cathodic-leading biphasic pulse, with 200µs per phase with no interleaving between pulses. The duration of the stimulus was adjusted to 500∼600ms to accommodate the imaging acquisition frequency. Throughout all recording sessions and across all subjects, the stimulation frequency was fixed at 200Hz.

Although similar, individual response thresholds slightly varied. For quantification, we defined the stimulation amplitude as Low, Medium, and High, as follows: at Low, mean=6.8µA with a standard deviation (SD) of ±1.1µA; Medium, mean=12.7, SD, ±1.2µA; and High, mean=15.9, SD, ±1.5µA. Electrode pairs were spaced between 370-850µm across experiments, depending on anatomical and surgical constraints. We then applied either parallel stimulation current intensities or with 1∼3µA difference to the other electrode, which was located out of two-photon recording field of view. Five to ten trials of stimulation were delivered for each condition, with a 10-second inter-trial interval. Between conditions, including single- and dual-electrode stimulation at various intensities, a 2∼3 minute inter-condition interval was applied.

### Data Analysis

#### Analysis of two-photon imaging data

For single-cell level analysis, we analyzed 6 mice for ‘medium’ stimulation condition, 5 mice for both ‘low’ and ‘high’ stimulation conditions with robust and stable fields of view throughout the experiment with visual inspections, ensuring all neuronal somata were clearly visible without response threshold changes or two-photon imaging plane shifts across single and dual-electrode stimulation conditions. 5 mice are shared for medium and high stimulation intensity analysis. We excluded the data from ‘low’ stimulation intensity (= the lowest stimulation intensity that was applied) from the in-depth statistical analysis because the number of responsive cells was too small for reliable statistics and quantification. We also excluded one mouse data for ‘high’ intensity stimulation analysis due to response threshold changes.

Signal processing and data analysis were all conducted using a customized Matlab pipeline. As the initial standardized preprocessing of the images, we used CalmAn (69) for motion correction, which utilizes the NoRMCorre algorithm(70) correcting non-rigid motion artifacts as our previous work (68). After the motion correction, we used Cellpose (71) (“cyto”) for cell segmentation, and then the extracted ROIs were manually corrected and added based on visual inspection. Then, neuropil contamination was subtracted with a coefficient of 0.4 using the fluorescence changes around the extracted soma ROIs (72).

### Statistical analysis

To determine the ROIs representing significantly responsive cells to micro-electrode stimulation, we selected cells whose mean fluorescence during the 2-second post-stimulation period exceeded three times the standard deviation (SD) of the mean baseline fluorescence measured from −3s to −1s before stimulation, using the trial averaged trace. The modulation magnitude (=modulatory effect), which can be expressed as coE1E2 – (sE1 + sE2), was determined by subtracting the cell’s response during single electrode from the response to dual electrode stimulation of. Baseline subtracted fluorescence was used within individual mouse statistics (ΔF), and pooling across all somata responses from all mice; the responses were normalized to the baseline within each mouse (ΔF/F0, normalized).

At the single-cell level, we performed a paired *t*-test for significance testing of modulation magnitude between dual and single electrode stimulation conditions across stimulation trials, and performed multiple comparison correction with α = 0.05 to control for false positives (73). Neuronal responses to single- and dual-electrode stimulation were then pooled across all animals for population-level analysis. Then, we used a *t*-test against zero for population-level statistics showing no difference between conditions across all neurons in the group of cells we categorized.

We used chi-square tests to assess the significance of differences in the proportion of cells exhibiting neuromodulation (either enhancement and attenuation) across different stimulation intensities.

To assess the significance of the correlation for each individual cell, we performed a bootstrapping procedure in which the pairings between response amplitudes in the single- and dual-electrode stimulation conditions were randomly shuffled (n=10000 iterations). For each iteration, the Pearson correlation coefficient was calculated, generating a null distribution of correlation values under the assumption of no relationship between conditions. The Monte Carlo *p*-value for each cell was then determined by comparing the observed correlation coefficient to this null distribution. At the population level, the significance of the mean correlation coefficient was tested using a *t*-test against zero, following Fisher-Z-transformation of individual cell correlation coefficients to normalize their distribution.

To visualize the relationship between variables and estimate confidence intervals for the fitted curve, we applied bootstrapping (n=2000 resamples with replacement) to the data points used for locally weighted scatterplot smoothing (LOWESS) fitting. Then, the 2.5^th^ and 97.5^th^ percentiles of the fitted values at each point along the x-axis were taken to represent the 95% confidence interval.

To assess the significance of the modulatory effect during dual-electrode stimulation as a function of single-electrode stimulation, we applied a sliding window approach with different parameters for responsive and non-responsive cells (=Non_Resp cells). For E1_Resp Cell, we used a window size of 50 bins with a step size of 20 bins, allowing for overlap. For Non_Resp cells, due to their smaller response amplitude range, we applied a sliding window of 15 bins with a step size of 5 bins. At each bin, a one-sample t-test was performed against zero, and the resulting p-values were corrected for multiple comparisons using False Discovery Rate (FDR) correction (73).

For testing of the significantly enhanced and significantly attenuated locations (=distance between soma and the electrodes) across the population of neurons across recording sites, we created a surrogate dataset (n=10000) by randomizing the distance and obtained a smoothed (”lowess”) fitting curve. Then, Monte-Carlo *p*-value (*p*<0.05) was estimated between the surrogate and the real dataset, which was further corrected with FDR. For comparing computational model scenarios, we used Wilcoxon Signed-Rank test for the significance testing between conditions.

### Connectivity network and computational modeling

To quantify the effect of neural network connectivity on modulation during dual-electrode stimulation, we modeled a population of neurons randomly positioned in space. Each neuron has a probability of connecting to other neurons based on their relative distance, defined by an exponential decay function,

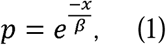

where *p* is the probability of connection, *x* is the distance between the two neurons, and *β* is the decay constant. This probability governs both synaptic connections between neurons and the connectivity between electrodes and neurons. For example, a probability of 10% implies that a presynaptic neuron is likely to connect to one postsynaptic neuron in a group of 10. The connection strength (i.e., weight) from each electrode to a neuron is also defined by Eq.1, such that neurons closer to an electrode receive a stronger input current.

In this model, electrode 1 (E1) is placed at the edge of the neuronal populations, while electrode 2 (E2) is positioned 485µm away, replicating the experimental inter-electrode spacing (370-858µm). Each electrode delivers pulses of 7680pA for a duration of 500ms. The effective current received by each neuron is scaled by its connection weight from the electrode. The stimulation protocol lasts 20 seconds per trial. After an initial current pulse to excite the system, the first two pulses are delivered solely by E1, followed by two pulses from E2, and concluding with two simultaneous pulses from both E1 and E2 (coE1E2). This sequence enables quantification of the modulatory effects of single versus dual-electrode stimulation.

Noise is introduced across the network with a mean of 0pA and a standard deviation equal to the pulse amplitude. Noise is sampled independently at every 0.1ms time step and applied to all neurons, allowing both positive and negative fluctuations. For each stimulation scenario, we generated 50 unique random seeds to ensure variability across trials. These seeds govern the neuron positions, synaptic connectivity and noise patterns. The same seed set is used across different test scenarios – e.g., removing inhibitory input or omitting noise input – allowing direct comparison under identical initial conditions.

A representative connectivity heatmap (Supplementary Figure 5) illustrates the spatial distribution of connection weights from each electrode and highlights neurons receiving input from both electrodes. The relevant model parameters, including decay constants are summarized in Table 1.

### Multi-layer perceptron regression for modulation prediction

#### Feature Engineering

We extracted 10 initial features that reflect the geometric and stimulation-related context of each neuron (Table 2). These include neuron-to-electrode distances and angles (for E1 and E2), distance between electrodes, prestimulation ΔF/F range, stimulation intensities from each electrode, and the electric field magnitude at the soma (Fig. 6C). The electric field was simulated in 3D using finite element modelling (FEM) in COMSOL Multiphysics and calculated as the vector norm of the field at each neuron location. To improve predictive performance, 7 additional interaction features were calculated, resulting in 17 total input features.

The response variable was defined as the mean area-under-the-curve (AUC) of the ΔF/F normalized calcium trace per neuron, averaged across stimulation peaks. AUC was computed using the trapezoidal rule over N+1 evenly spaced points and normalized by the imaging frame rates. To stabilize variance and reduce skewness, response values were transformed using the Box-Cox power transformation.

#### Model setup

We used a Multi-Layer Perceptron (MLP) Regressor to predict neuronal responses based on the extracted features, written in Python using Sci-Kit Learn (74) and own code. We used a dataset of 994 neurons recorded across 10 stimulation conditions (N=5 mice). One mouse was excluded from this analysis due to a limited number of neurons, resulting from a smaller two-photon field of view.

An MLP Regressor is a type of artificial neural network optimized for continuous output prediction. It consists of an input layer, multiple hidden layers, and an output layer. Each neuron applies a weighted sum of its inputs, followed by a nonlinear activation function, enabling the network to learn complex, nonlinear mappings. The model was trained via backpropagation using gradient descent to minimize mean squared error (MSE) loss.

The neural network consisted of 3 hidden layers of 100 units each and a bias term (Figure 6E), ReLU activation function, an L2 regularization (α = 0.0001), and the Adam optimizer. Input features were min-max normalized using parameters fit on the training set, and the same transformation was applied to both train and test sets. Samples were randomly shuffled before training.

#### Generation and simulation of artificial neuronal populations

We trained a complementary single-electrode model using data from single-electrode stimulation to predict neuronal responses under these conditions. To extend our analysis on neuromodulation effect from the model, we generated a simulated population of 400 neurons evenly distributed within a 350 x 350 μm field of view (20 neurons per row, 17.5μm apart). Electrodes were placed 400μm apart, near opposite edges of the field of view. The simulation field was expanded to include both electrodes, enabling the visualization of predicted responses beyond the empirical imaging area. Model input features were computed as in the empirical dataset, with pre-stimulation range values randomly sampled from the empirical distribution. After min-max scaling, neuronal responses were predicted at each coordinate using the trained MLP Regressor, and predicted responses were reverse-Box Cox transformed to obtain ΔF/F values.

To quantify the modulation effect, we subtracted the simulated single-electrode responses (E1 and E2) from the coE1E2 response and applied a LOWESS curve fit. Simulation accuracy was validated based on the performance of the trained MLP Regressor model (see Methods, MLP regressor).

---

**S1.**
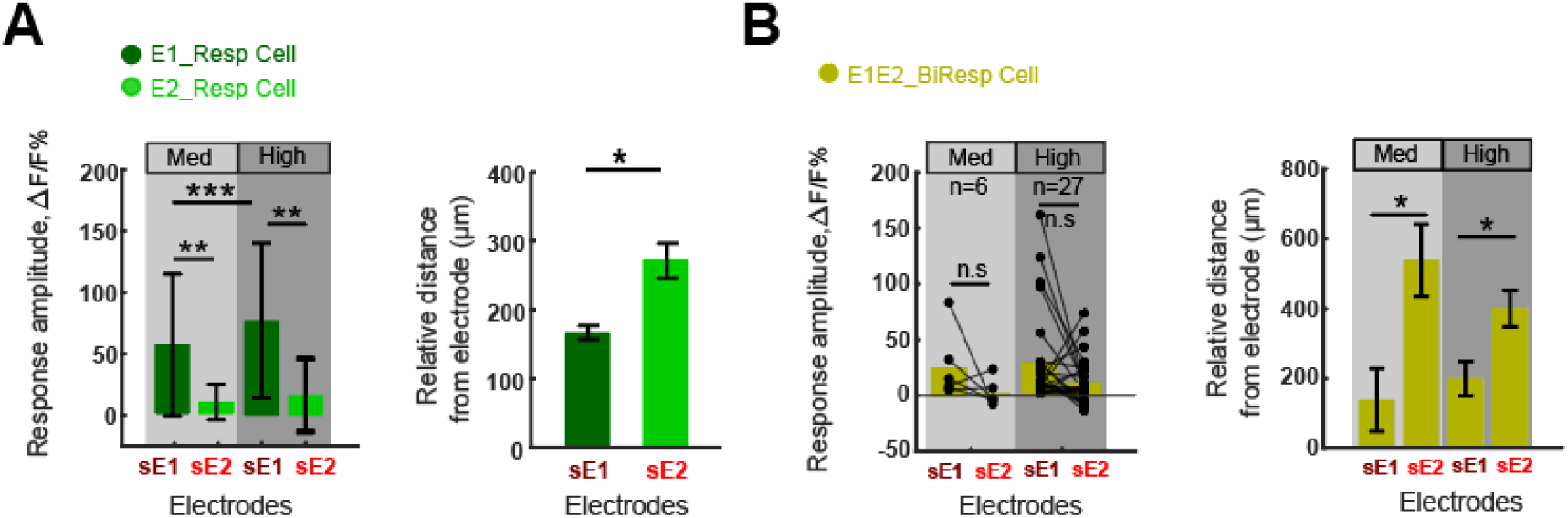
Response amplitude and soma distance from electrodes during single-electrode stimulation. **A.** Response amplitude (left), relative distance of soma from electrode (right) comparison E1_Resp Cell (dark green), E2_Resp Cell (green). **B**. Same convention as A, but showing Bimodal cells (\**p*<0.05, \*\**p*<0.005, *** *p*<0.0001).

**S2.**
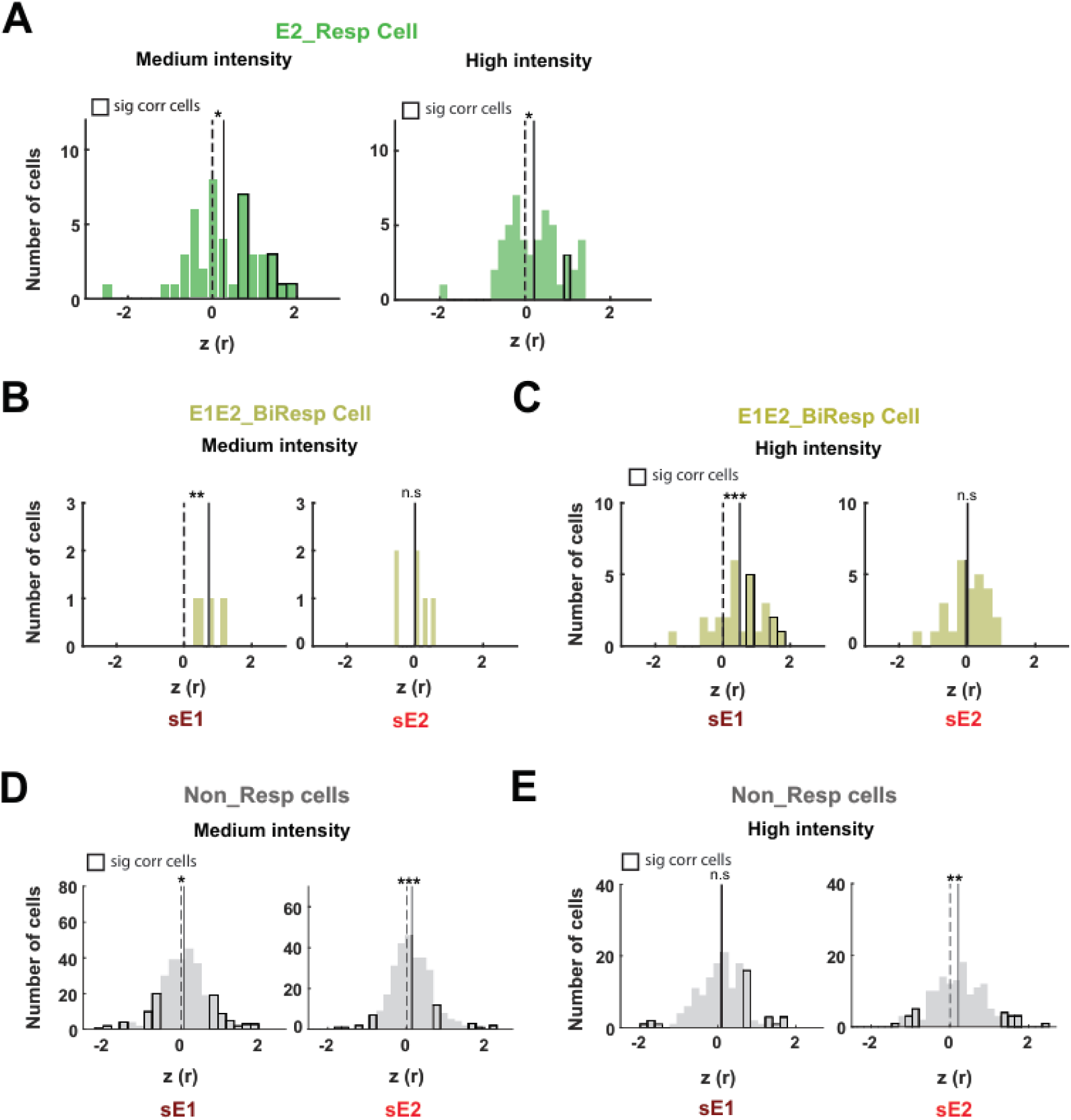
Population-level cell correlation is biased to positive correlation. **A**. E2_Resp cell at medium (left) and high (right) stimulation intensity. Black edge color bar indicates cells showing significant response amplitude correlations between single and double conditions. The dashed line is at zero; the solid line indicates the mean correlation coefficient, Fisher-z transformed for normality. **B-E**. Same convention as A, but for bimodal cells (**B,C**) at medium intensity (**B**) showing the correlation between double and single sE1 (left) and single sE2 (right), and at high intensity (**C**). Non_Resp cells are shown in **D,E**. (\**p*<0.05, \*\**p*<0.01, \*\*\**p*<0.005).

**S3.**
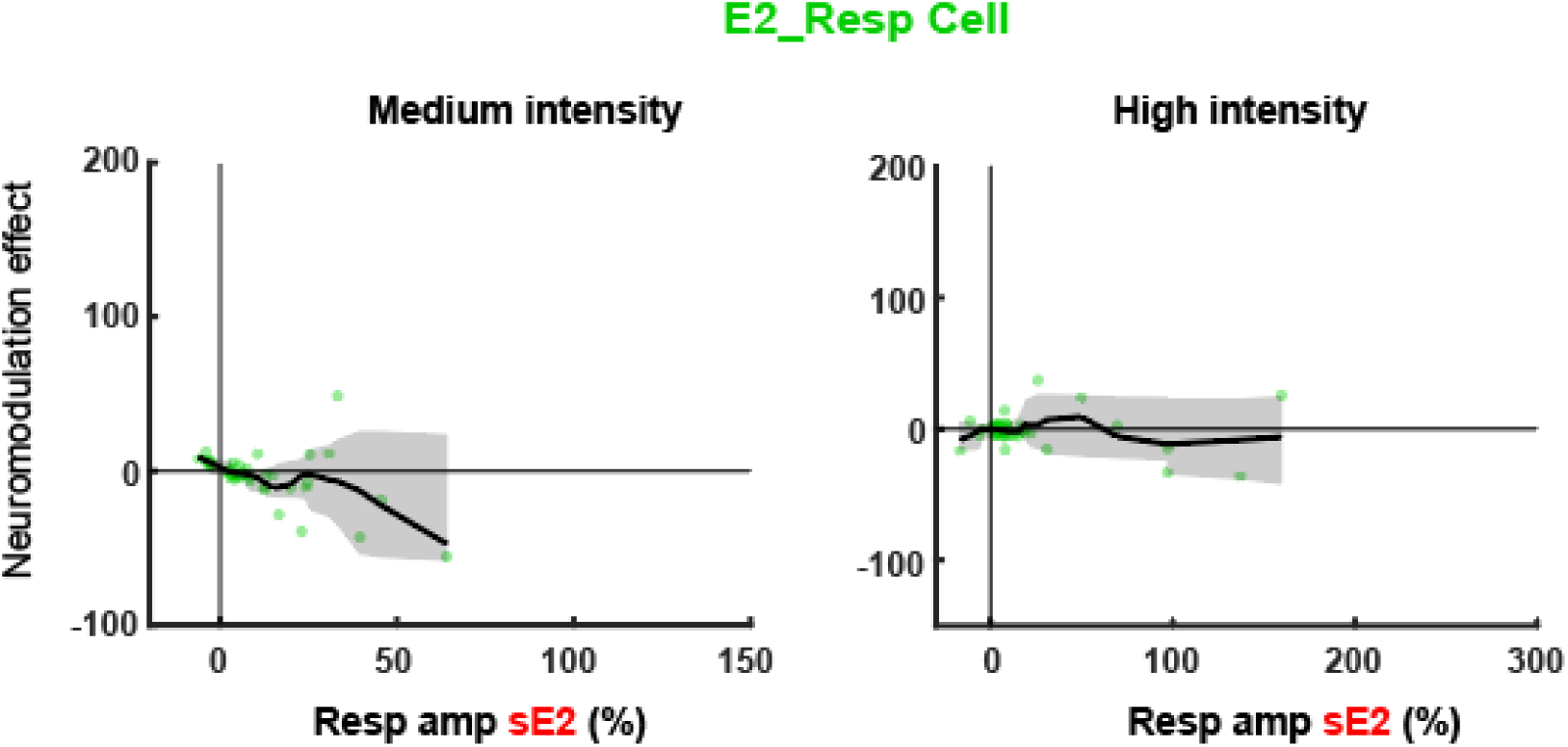
Same convention as Figure 3E-F, but showing neuromodulation effect (%) as a function of somata response among E2_Resp cell.

**S4.**
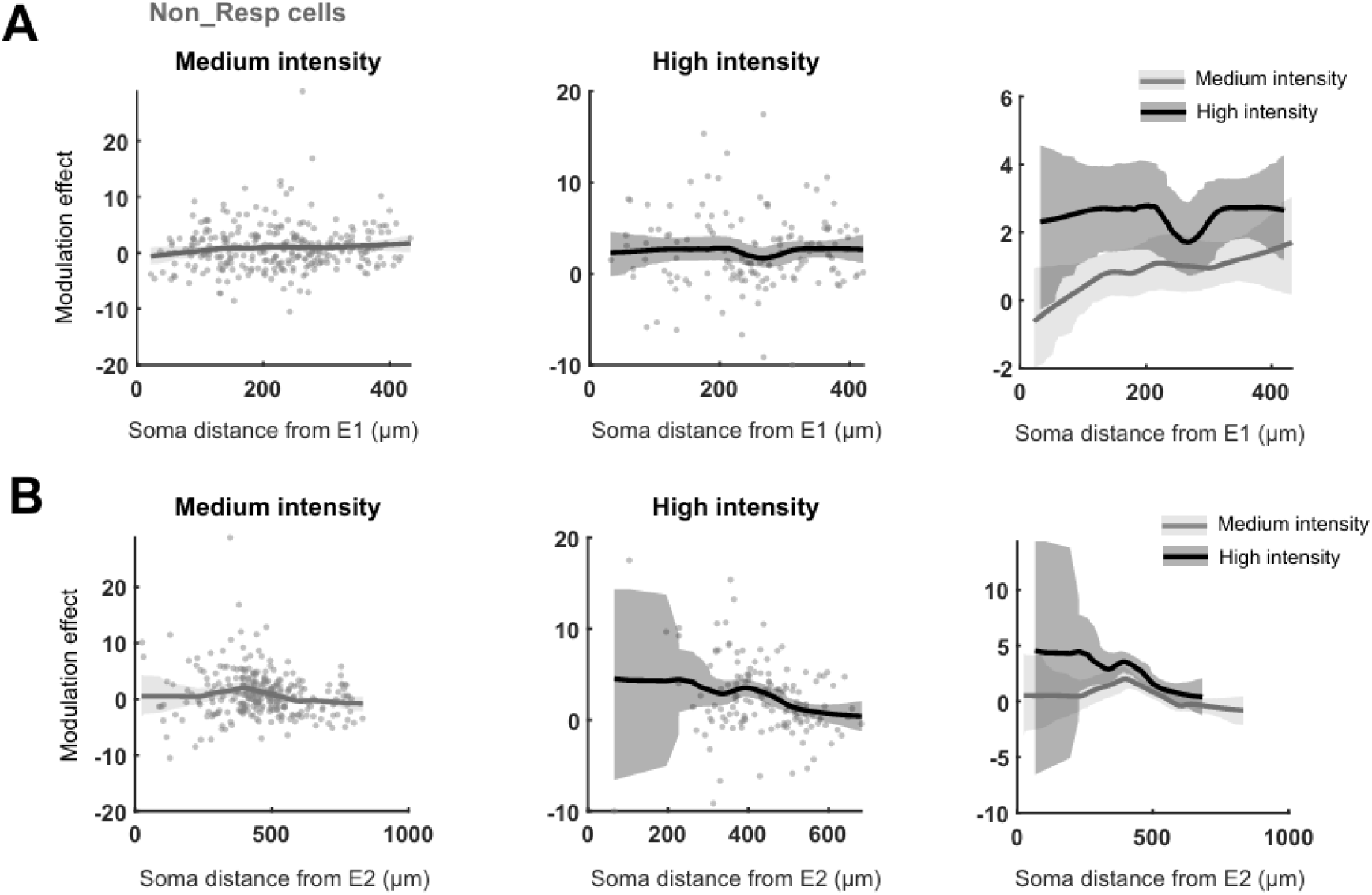
No diverging modulation effect shown between different current intensities among Non_Resp cells. **A.** Response amplitude (y-axis), relative distance of soma from E1 (x-axis). Medium (left), high (middle) stimulation intensities. Overlaid data between two conditions (right), medium (black), high (gray) intensities. **B**. Same convention as A, but somata locations are relative to E2. Black contour denotes LOWESS fit to data with shade of 95% confidence interval obtained by bootstrapping.

**S5.**
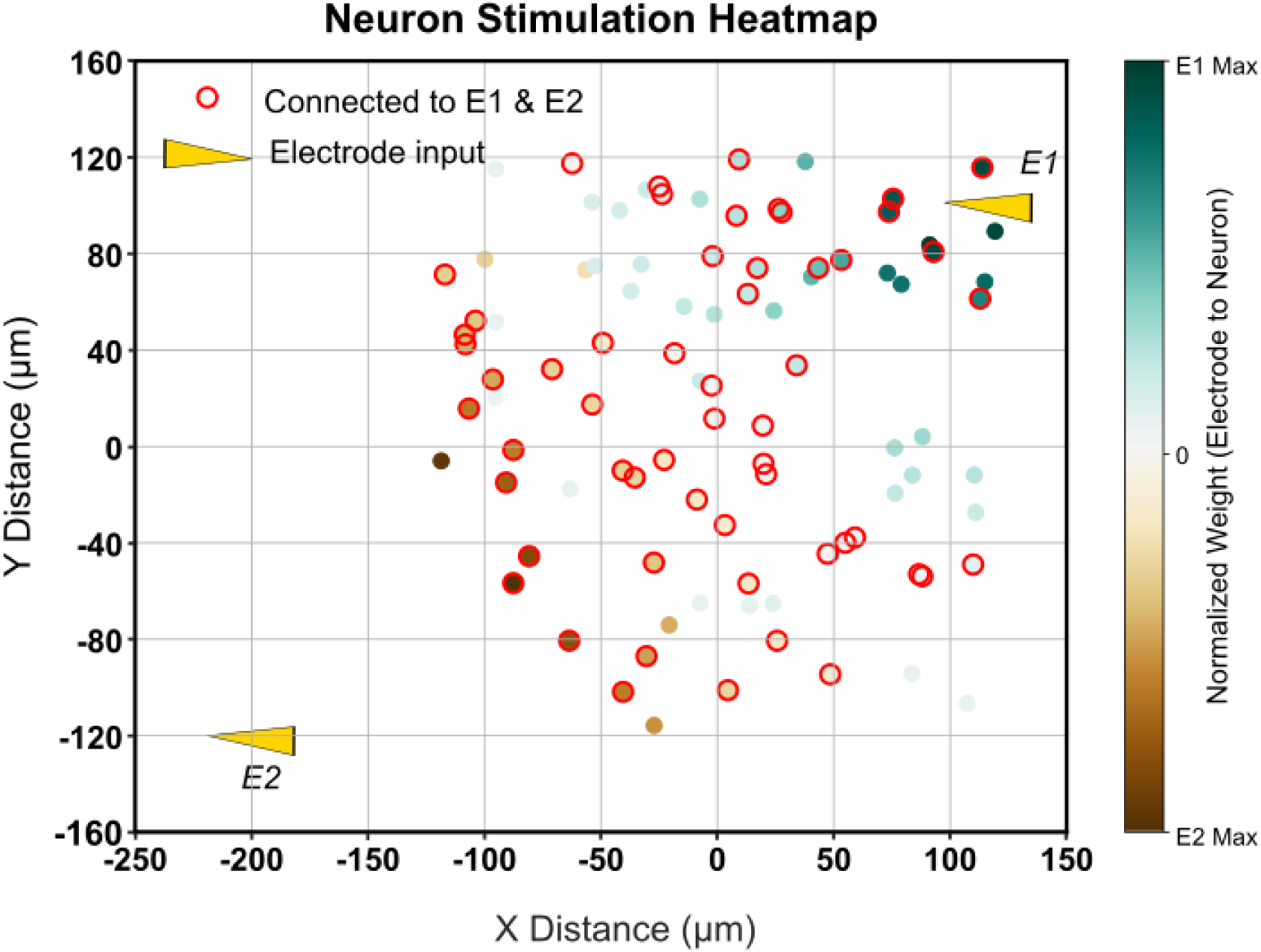
Electrode connectivity heatmap. Neurons shown in green receive input from E1, while brown represents E2. Neurons circled in red receive input from both electrodes and the fill color of the dot represents the electrode with a stronger connection.

**S6.**
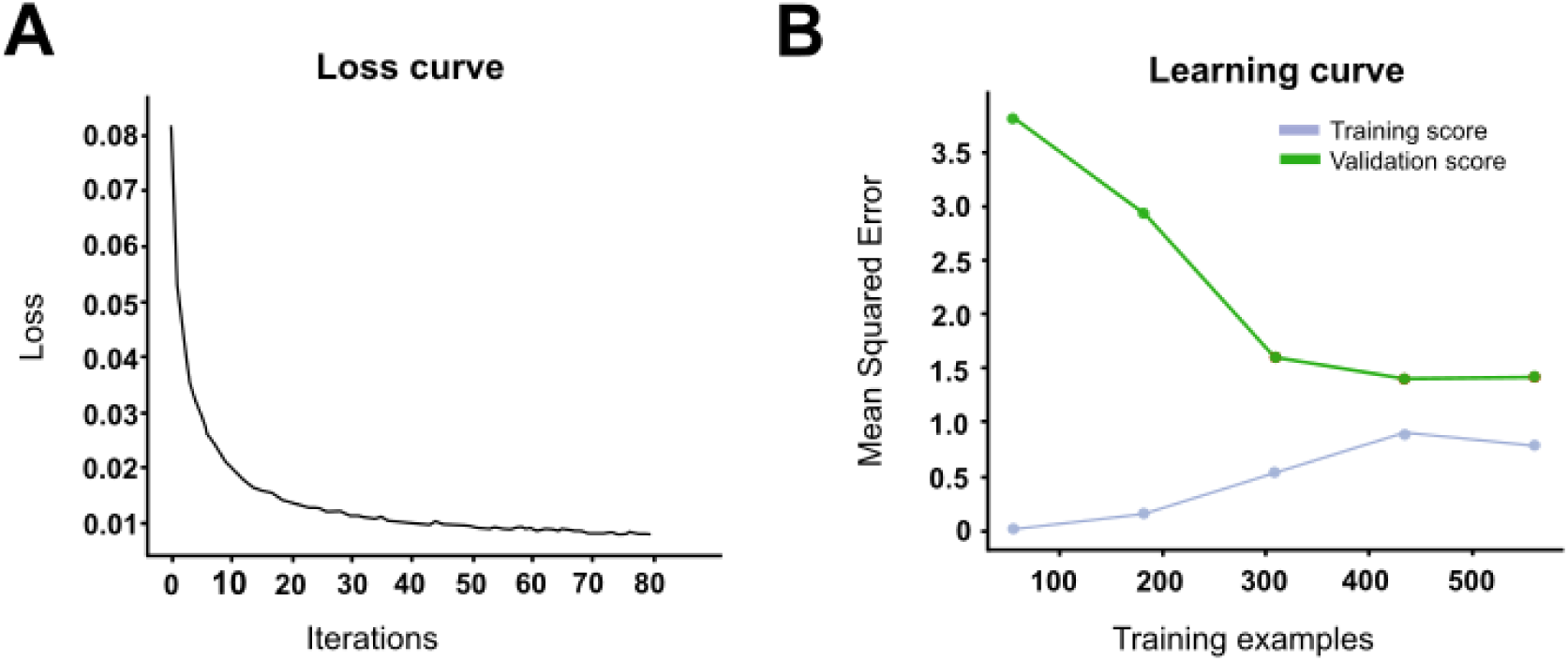
Model architecture and performance validation. **A**. Training loss curve showing decreasing loss, indicating good fit. **B**. Learning curve with MSE. Training score (gray) and validation score (green).

